# Wilcoxon rank-sum test still outperforms dearseq after accounting for the normalization impact in semi-synthetic RNA-seq data simulation

**DOI:** 10.1101/2022.06.07.494963

**Authors:** Yumei Li, Xinzhou Ge, Fanglue Peng, Wei Li, Jingyi Jessica Li

**Affiliations:** Division of Computational Biomedicine, Department of Biological Chemistry, School of Medicine, University of California, Irvine, Irvine, CA 92697, USA; Department of Statistics, University of California, Los Angeles, CA 90095; Department of Molecular and Cellular Biology, Baylor College of Medicine, Houston, TX 77030, USA; Interdepartmental Program in Bioinformatics, University of California, Los Angeles, CA 90095; Department of Human Genetics, University of California, Los Angeles, CA 90095; Department of Computational Medicine, University of California, Los Angeles, CA 90095; Department of Biostatistics, University of California, Los Angeles, CA 90095

## Abstract

In this response to the correspondence by Hejblum et al. [1], we clarify the reasons why we ran the Wilcoxon rank-sum test on the semi-synthetic RNA-seq samples without normalization, and why we could only run dearseq with its built-in normalization, in our published study [2]. We also argue that no normalization should be performed on the semi-synthetic samples. Hence, for a fairer method comparison and using the updated dearseq package by Hejblum et al., we re-run the six differential expression methods (DESeq2, edgeR, limma-voom, dearseq, NOISeq, and the Wilcoxon rank-sum test) without normalizing the semi-synthetic samples, i.e., under the “No normalization” scheme in [1]. Our updated results show that the Wilcoxon rank-sum test is still the best method in terms of false discovery rate (FDR) control and power performance under all settings investigated.

## Key messages in Hejblum et al. [1]

The correspondence by Hejblum et al. [1] pointed out the effect of normalization on our false discovery rate (FDR) benchmarking result of differential expression (DE) methods using semi-synthetic data [2] (Figure 2 and Figures S20-S30 in [2]). In particular, when we ran the Wilcoxon rank-sum test on permuted semi-synthetic samples to verify FDR control, we did not include a normalization step; instead, when we ran the other five DE gene methods (DESeq2, edgeR, limma-voom, NOISeq, and dearseq), we included their built-in normalization. Hence, our comparison results (Figure 2 and Figures S20-S30 in [2]) did not put the Wilcoxon rank-sum test on the same ground as the other five DE methods (in terms of the use of normalization). Hejblum et al. showed that, if the Wilcoxon rank-sum test was run on normalized semi-synthetic samples, it would also have inflated FDR as the other five DE methods did.

Another key message in Hejblum et al. [1] is that benchmarking DE methods using semi-synthetic samples has three schemes: (1) “Permutation first,” (2) “No normalization,” and (3) “Normalization first.” Our previous benchmarking used scheme (1) for DESeq2, edgeR, limma-voom, NOISeq, and dearseq, but scheme (2) for the Wilcoxon rank-sum test [2] (Figure 2 and Figures S20-S30 in [2]). Unlike schemes (1) and (2), scheme (3) requires a different way of generating semi-synthetic samples (Table 1; detailed in the next two paragraphs). Hejblum et al. re-did the benchmarking under each scheme and argued that dearseq [3] outperforms the other five methods, including the Wilcoxon rank-sum test.

**Table 1.**
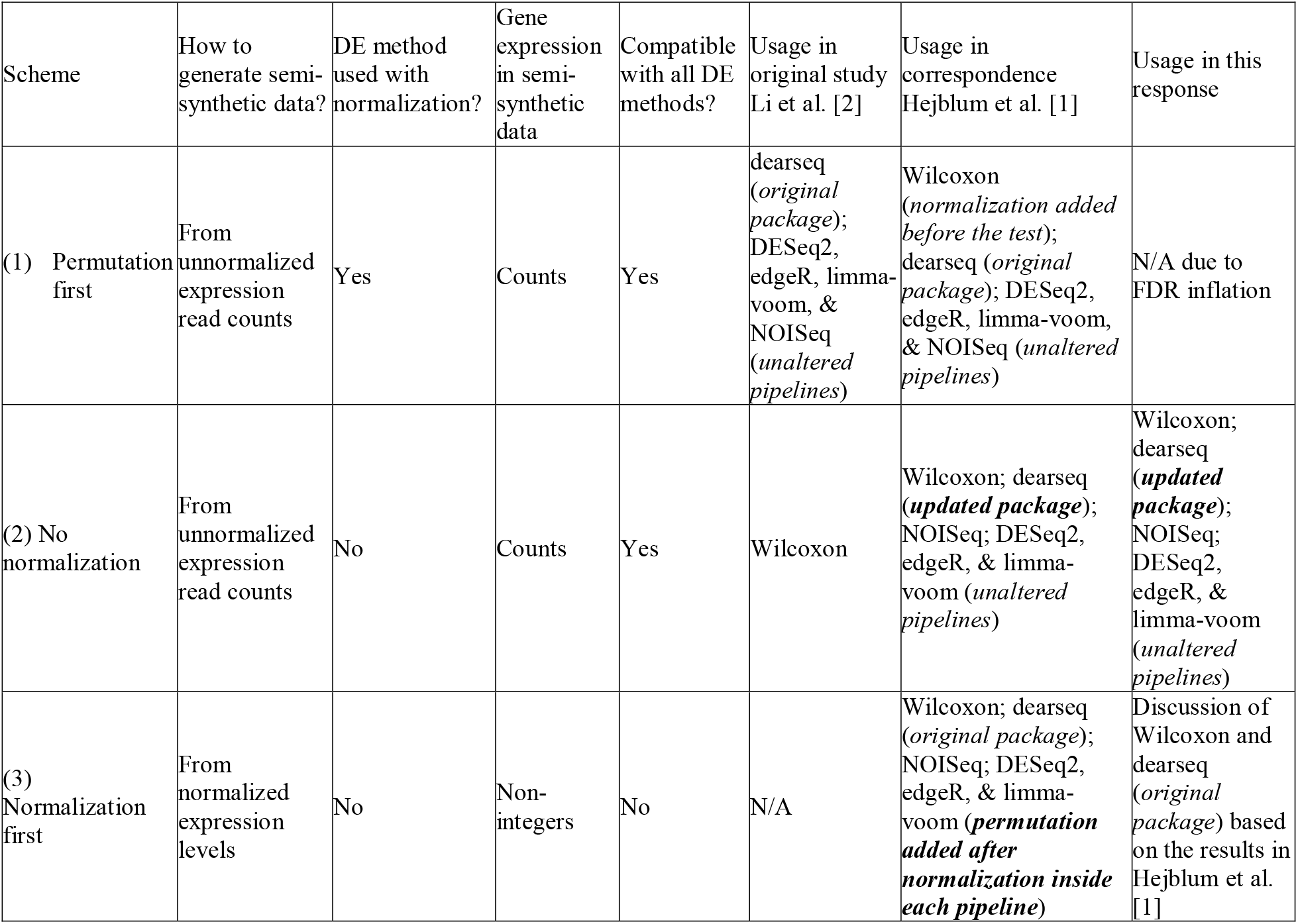
Properties and usages of the three (semi-synthetic data generation + DE method implementation) schemes proposed in Hejblum et al. [1]

### Clarification of the three (semi-synthetic data generation + DE method implementation) schemes proposed in Hejblum et al. [1]

In our published study [1], we generated semi-synthetic samples by defining the true differentially expressed genes (DEGs) as those identified by all six DE methods at a very small FDR threshold (0.0001%). For these true DEGs, we fixed their unnormalized expression read counts in the real data’s two conditions. For the other genes, we considered them as true non-DEGs and randomly permuted their unnormalized expression read counts between the real data’s two conditions. Because of random permutation, in the semi-synthetic data, each true non-DEG’s counts under the two conditions are from the same distribution, so no normalization is needed, and scheme (2) should be used. Hejblum et al. [1] pointed out that post-permutation normalization, i.e., scheme (1), would distort each true non-DEG’s expression levels under the two conditions (because true DEG’s different expression counts in different samples would make the samples have different library sizes) and cause true non-DEGs to be identified as false positives, a point we completely agree.

Besides, Hejblum et al. [1] proposed scheme (3) that generates semi-synthetic samples in a different way: real data are first normalized within each condition, and then semi-synthetic data are generated by fixing true DEGs’ normalized expression levels in the real data’s two conditions and randomly permuting true non-DEG’s normalized expression levels between the real data’s two conditions. It is true that this scheme (3), like scheme (2), does not perform post-permutation normalization, so true non-DEG’s expression levels would not be distorted and lead to false positives. However, we argue that scheme (3) would generate semi-synthetic samples containing normalized expression levels that are no longer counts, making many DE methods that require count data input become inapplicable. In fact, Hejblum et al. [1] had to alter each DE method pipeline by adding a permutation step after the normalization step, making scheme (3) not only a semi-synthetic data generation scheme but also an altered implementation of DE methods.

Because of the FDR inflation caused by scheme (1) and the complexity of using scheme (3), our recommendation is scheme (2).

### Explanation of why the Wilcoxon rank-sum test was run under scheme (2) but dearseq could only be run under scheme (1)

We agree with Hejblum et al. that scheme (1) would inflate the FDR. In fact, this is the reason why we used scheme (2) for the Wilcoxon rank-sum test, which is not an RNA-seq-specific software package and thus does not include a built-in normalization step. However, the other five RNA-seq-specific DE methods include built-in normalization, and users would likely use them as pipelines by not removing the built-in normalization step. Hence, while we agree that running them as pipelines, i.e., under scheme (1), was not fair for evaluating the FDR control of their statistical tests, we believe that our previous results (Figure 2 and Figures S20-S30 in [2]), in conjunction with Hejblum et al.’s results, are meaningful for showing the risks of using bioinformatics tools as black-box pipelines.

Moreover, we would like to point out that the dearseq R package we used in [2] (Bioconductor date Oct 26, 2021) does not support running dearseq under scheme (2). Hence, running dearseq under scheme (1) was our only option given that we would like to use count data as input for all DE methods. We have a detailed discussion on this issue in **Additional file 1**.

### The Wilcoxon rank-sum test still outperforms dearseq under schemes (2) and (3)

While we agree that scheme (1) does not provide a fair FDR evaluation because of the bias introduced by post-permutation normalization, we do not agree with Hejblum et al. [1] that dearseq outperforms the Wilcoxon rank-sum test under schemes (2) and (3). Note that dearseq has two versions: dearseq (permutation) uses a permutation test for p-value calculation, while dearseq (asymptotic) uses an asymptotic test for p-value calculation.

Under scheme (2), we re-run the six DE methods and obtained results similar to those in Hejblum et al. [1]. Our **Figs. 1 and 2** and Hejblum et al.’s Figure 1 [1] show that dearseq (permutation) cannot control the FDR, and dearseq (asymptotic) can control the FDR only when the sample size is large enough (e.g., sample size ∼40 when the target FDR is 0.1%; **Fig. 2**). In contrast, the Wilcoxon rank-sum test has consistent FDR control across all sample sizes. **Fig. 1** and **Fig. 2** also show that under the same actual FDR, dearseq (permutation) has the worst power among all DE methods, while dearseq (asymptotic) has no obvious power advantage over the Wilcoxon rank-sum test. Based on these results, we argue that the Wilcoxon rank-sum test outperforms dearseq (asymptotic), and obviously dearseq (permutation), in terms of FDR control consistency and power.

**Fig. 1.**
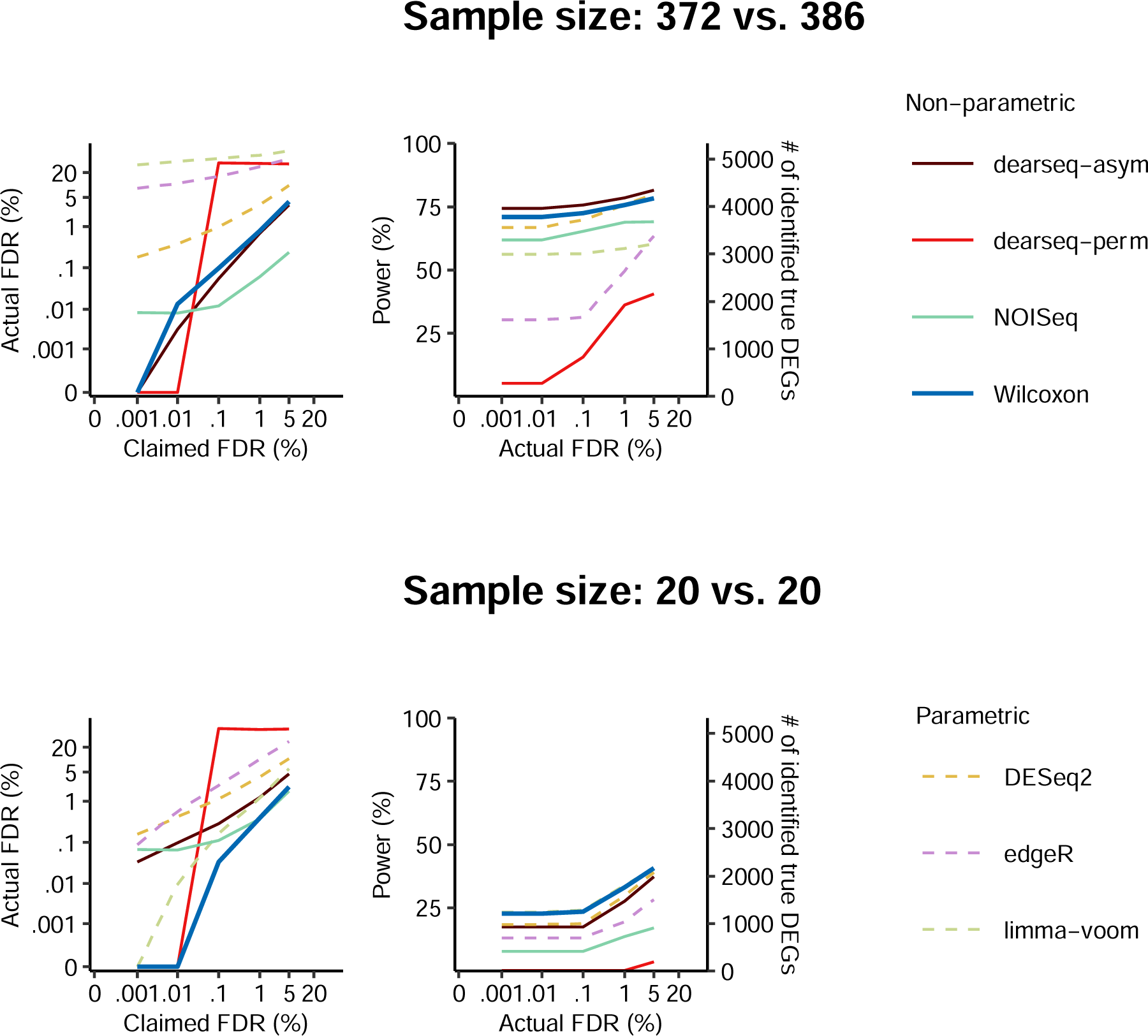
The comparison of DE methods on heart left ventricle vs. atrial appendage GTEx datasets with semi-synthetic ground truths under the No normalization data generation scheme. The FDR control (left panel), and power given the actual FDRs (right panel) under a range of FDR thresholds from 0.001% to 5% for sample sizes: all sample from the two condition (top) and 20 samples per condition (bottom). The Wilcoxon rank-sum test control the FDR and achieve good power under all FDR thresholds for both sample sizes.

**Fig. 2.**
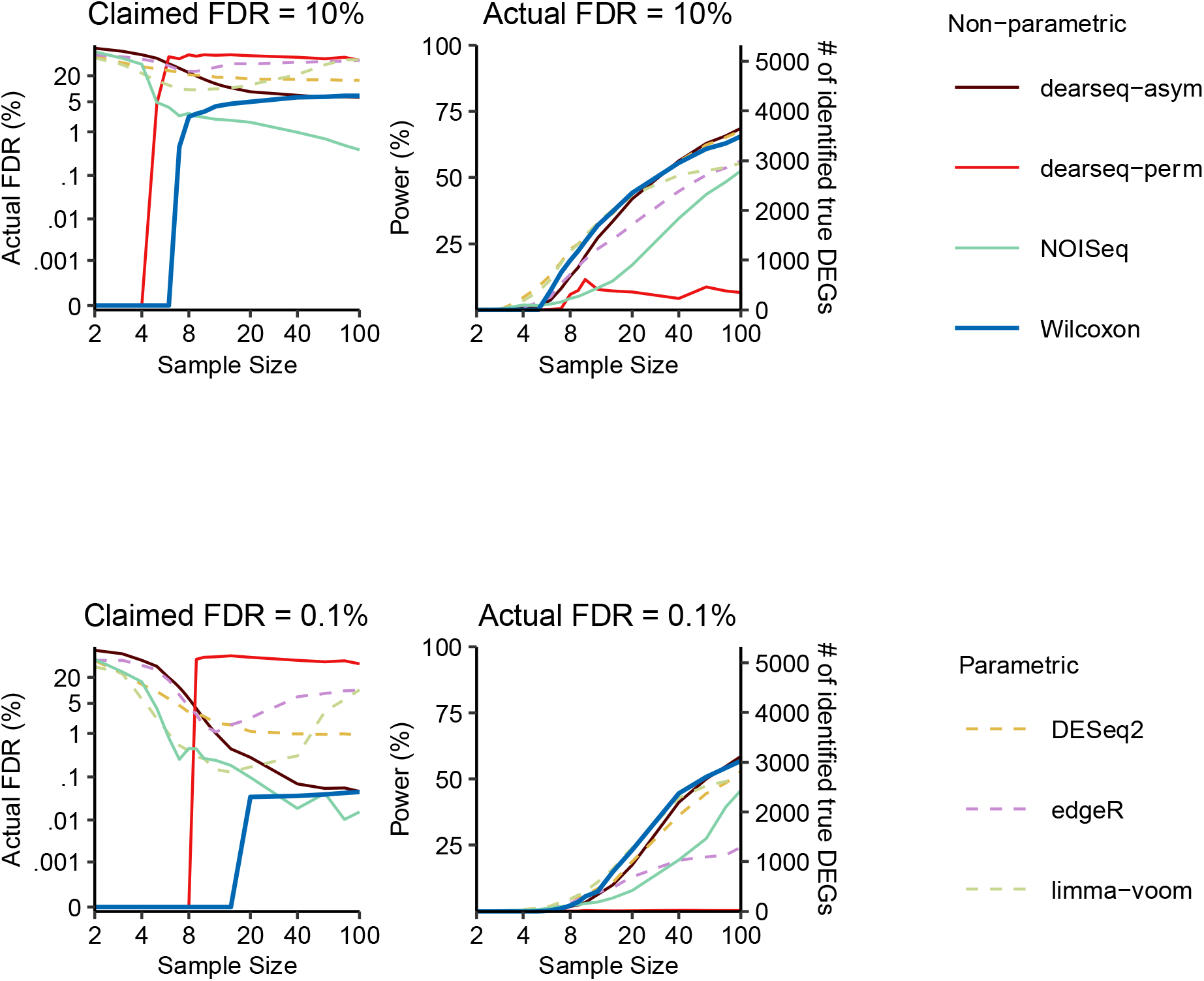
The comparison of DE methods on heart left ventricle vs. atrial appendage GTEx datasets with semi-synthetic ground truths under the No normalization data generation scheme. The FDR control (left), and power given the actual FDRs (right) for a range of per-condition sample sizes from 2 to 100, under FDR thresholds 10% (top panels) and 0.1% (bottom panels). The claimed FDRs, actual FDRs, and power were all calculated as the averages of 50 randomly down-sampled datasets.

Since scheme (3) is not directly applicable to DESeq2 [4], edgeR [5], and limma-voom [6], which only accept gene expression read counts as input data, we choose not to alter their pipelines and run them under this scheme. Hence, we use the results in Hejblum et al. [1] to compare the Wilcoxon rank-sum test with dearseq (permutation) and dearseq (asymptotics). Hejblum et al.’s Figures 1 and 2 show that dearseq (permutation) can control the FDR but lacks power, and dearseq (asymptotics) can control the FDR only when the sample size is large enough. In contrast, the Wilcoxon rank-sum test has consistent FDR control across all sample sizes. Moreover, dearseq (asymptotics) does not have an obvious power advantage over the Wilcoxon rank-sum test. Hence, we argue that the Wilcoxon rank-sum test is also the preferred choice under scheme (3).

Although we do not agree with Hejblum et al. that dearseq outperforms the Wilcoxon rank-sum test in terms of FDR control and power for two-condition comparisons, we agree that dearseq can account for more complex experimental designs.

### Final note

We would like to clarify that our study [2] was not a comprehensive benchmark. Many DE methods have been developed in the last decade (including more than 20 methods that have been benchmarked in previous studies; see Table S1 in [1]), and it is possible that some methods may outperform the Wilcoxon rank-sum test on specific datasets. Our study [2] aimed to emphasize the importance of sanity check and voice the cautionary message that using popular methods such as DESeq2 [4] and edgeR [5] blindly may lead to excessive false positives.

## Availability of data and materials

All the codes and data used to generate the new results can be found at **Data and codes_response.zip** from Zenodo: https://doi.org/10.5281/zenodo.5241320.

## Ethics approval and consent to participate

Not applicable.

## Consent for publication

Not applicable

## Author contributions

X.Z. performed the data analysis. Y.L., X.Z., J.J.L and W.L. wrote the manuscript.

## Acknowledgements

We thank other members of Wei Li lab and Jingyi Jessica Li lab for helpful discussions.

## Supplementary information

Additional file 1: Document showing the dearseq version issue

## References

1. Hejblum BP, Ba K, Thibaut R, Agniel D: Neglecting normalization impact in semi-synthetic RNA-seq data simulation generates artificial false positives. bioRxiv 2022.

2. Li Y, Ge X, Peng F, Li W, Li JJ: Exaggerated false positives by popular differential expression methods when analyzing human population samples. Genome biology 2022, 23:1--13.

3. Gauthier M, Agniel D, Thibaut R, Hejblum BP: dearseq: a variance component score test for RNA-Seq differential analysis that effectively controls the false discovery rate. NAR genomics and bioinformatics 2020, 2:lqaa093.

4. Love MI, Huber W, Anders S: Moderated estimation of fold change and dispersion for RNA-seq data with DESeq2. Genome biology 2014, 15:1--21.

5. Robinson MD, McCarthy DJ, Smyth GK: edgeR: a Bioconductor package for differential expression analysis of digital gene expression data. Bioinformatics 2010, 26:139--140.

6. Law CW, Chen Y, Shi W, Smyth GK: voom: Precision weights unlock linear model analysis tools for RNA-seq read counts. Genome biology 2014, 15:1--17.

