## Additional file 1 for "Wilcoxon rank-sum test still outperforms dearseq after accounting for the normalization impact in semi-synthetic RNA-seq data simulation"

### dearseq version 1.6.0 does not support analysis under "No normalization" scheme

Xinzhou Ge

5/17/2022

In our paper "Exaggerated false positives by popular differential expression methods when analyzing human population samples," we used synthetic data analysis to show that existing DEG identification tools cannot control the FDR. Hejblum et al. recently submitted correspondence to our paper, which mentioned that we did not make a fair comparison in our benchmark analysis using semi-synthetic dataset. In particular, when we ran the Wilcoxon rank-sum test on permuted semi-synthetic samples to verify FDR control, we did not include a normalization step; instead, when we ran dearseq, we included the normalization step. They modified the semi-synthetic analysis, and argued that dearseq has good performance if it is used without the normalization step. However, we find that their results are based on the new version of dearseq, which was unavailable when we published our paper (Mar 15, 2022). This document shows that dearseq version 1.6.0, the latest version when our paper was published, does not support analysis under "No normalization" scheme.

We first install version 1.6.0 of dearseq. This version was released on Bioconductor on Oct 26, 2021, and it was the latest version when our paper was published. The source file can be found in http://bioconductor.org/packages/3.14/bioc/html/dearseq.html.

```
install.packages("dearseq_1.6.0.tar.gz",repos=NULL, type="source")
library(dearseq)
library(matrixStats)
```

We then try to repeat the analysis in their correspondence. Here we use their codes in zenodo. For simplicity, below, we only show the commands about their benchmark analysis under the no-normalization setting (codes are from Simulations/dearseqAsymp__3norms.R in their [zenodo website] (https://zenodo.org/record/6514317#.YoPbIy1h1bU)).

```
count_raw <- read.table("HeartAtrialAppendage.HeartLeftVentricle.gene_readCount.tsv",
                        sep = '\t', header = TRUE, row.names = 1)
rownames(count_raw) <- as.character(1:nrow(count_raw))
conditions <- read.table("HeartAtrialAppendage.HeartLeftVentricle.conditions.tsv",
                         sep = '\t')
condition_fact <- as.factor(as.matrix(conditions)[1, ])
dis0 <- readRDS("HeartAtrialAppendage.HeartLeftVentricle_dis.rds")
trueDE <- Reduce(intersect, dis0)
# Only look at the "No normalization" scheme
norm = "nonorm"
readCount <- count_raw
condition <- condition_fact
sampsize <- "full"

if(norm =="permprenorm"){
  # Permutation under H0
  trueDE_samplehalf <- sample(trueDE, length(trueDE)/2)
  readCount[!(rownames(readCount) %in% trueDE_samplehalf), ] <-
```

```r
    t(apply(readCount[!(rownames(readCount) %in% trueDE_samplehalf), ], 1, sample))
}

if(norm !="nonorm"){
  # Normalization
  y <- DGEList(counts=readCount, group=condition)
  keep <- filterByExpr(y)
  y <- y[keep, keep.lib.sizes=FALSE]
  y <- calcNormFactors(y,method="TMM")
  count_norm <- cpm(y, log=TRUE)
}else{
  count_norm <- as.matrix(readCount)
}

if(norm == "permpostnorm" | norm == "nonorm"){
  # Permutation under H0
  trueDE_samplehalf <- sample(trueDE, length(trueDE)/2)
  count_norm[!(rownames(count_norm) %in% trueDE_samplehalf), ] <-
    t(apply(count_norm[!(rownames(count_norm) %in% trueDE_samplehalf), ], 1, sample))
}

# Remove zero variance genes
count_norm <- count_norm[matrixStats::rowVars(count_norm)>0, ]

# Estimation
conditions_mat <- model.matrix(~ 1 + condition)[, 2, drop=FALSE]
```

Then we will have errors when we run the following command, :

```r
dearseq_asymp_res_pvals <- dearseq::dear_seq(exprmat=count_norm,
                                             variables2test=conditions_mat,
                                             which_test="asymptotic",
                                             parallel_comp=FALSE,
                                             preprocessed=T,
                                             which_weights = "loclin")$pvals
```

```
## Error in rlbin(x, y, gpoints, truncate): NA/NaN/Inf in foreign function call (arg 2)
```

This error disappears when we change the preprocessed argument into `preprocessed=F`. However, it means that the input matrix is not preprocessed, and in this case, a build-in normalization will be performed. To summarize, there is no way to correctly run dearseq on the permuted dataset without doing any normalization.

However, when we installed the latest version, version 1.8.1, which was released on Bioconductor on Apr 28, 2022, we did not see this error anymore.

```r
detach("package:dearseq", unload=TRUE)
install.packages("dearseq_1.8.1.tar.gz",repos=NULL, type="source")

library(dearseq)
dearseq_asymp_res_pvals <- dearseq::dear_seq(exprmat=count_norm,
                                             variables2test=conditions_mat,
                                             which_test="asymptotic",
                                             parallel_comp=FALSE,
                                             preprocessed=T,
                                             which_weights = "loclin")$pvals
```
